# Colocalization of dopamine D1 and D2 receptors with neuronal nitric oxide synthase in the rat paraventricular nucleus: a structural basis for dopaminergic–nitrergic interaction

**DOI:** 10.64898/2026.04.27.720621

**Authors:** Aixa R. Bello, Manuel Mas, Ricardo Reyes

## Abstract

The paraventricular nucleus (PVN) plays a central role in neuroendocrine and autonomic regulation, including male sexual behavior. Dopaminergic and nitrergic signaling within the PVN are functionally linked, but their cellular relationship remains unclear. We examined the colocalization of dopamine D1 and D2 receptors with neuronal nitric oxide synthase (nNOS) in adult male rats using double-label immunohistochemistry. nNOS-immunoreactive neurons were widely distributed in PVN, with subsets co-expressing D1R or D2R. Approximately 45% of nNOS-positive neurons expressed dopaminergic receptors. These findings provide structural evidence for dopaminergic–nitrergic interaction in the PVN and support the possibility of direct dopaminergic modulation of nitrergic neurons.

## Introduction

The paraventricular nucleus (PVN) of the hypothalamus is a major integrative center involved in the regulation of neuroendocrine, autonomic, and behavioral functions. Among its multiple roles, the PVN plays a critical part in the central control of male sexual behavior and penile erection (Argiolas and Melis, 2005).

Nitric oxide (NO), produced by neuronal nitric oxide synthase (nNOS), is a key neuromodulator in the central regulation of erectile function. In the PVN, activation of nitrergic neurons leads to NO release, which is essential for the initiation of penile erection through neuroendocrine and descending autonomic pathways (Chen and Chang, 2002; Ferrini et al., 2003; Melis and Argiolas, 2021; Melis et al., 2022).

Dopamine is another major neurotransmitter involved in the facilitation of sexual behavior. Dopaminergic stimulation within the PVN has been shown to induce penile erection, an effect associated with activation of nitric oxide production (Melis et al., 1996; Melis et al., 2000; Sanna et al., 2012; Melis et al., 2022). Both D1-like and D2-like receptors contribute to these effects; however, their precise cellular localization in relation to nitrergic neurons within the PVN remains poorly defined.

Thus, despite strong functional evidence supporting an interaction between dopaminergic and nitrergic systems, the structural basis underlying this relationship remains unclear. To our knowledge, this study provides the first direct anatomical evidence of colocalization between D1 and D2 dopamine receptors with nNOS in PVN neurons. Therefore, the aim of the present study was to investigate this colocalization in adult male rats, in order to provide an anatomical framework for understanding the neurochemical integration between these systems.

## Materials and methods

### Animals and tissue preparation

Adult male Sprague–Dawley rats (200–220 g) (n=6) were used in this study. All experimental procedures were conducted in accordance with European guidelines (Directive 2010/63/EU) and were approved by the Institutional Animal Care and Use Committee of the University of La Laguna (CEIBA2026-3812).

Animals were deeply anesthetized with sodium pentobarbital (120 mg/kg) and transcardially perfused with 0.1 M phosphate buffer (pH 7.4) followed by 4% paraformaldehyde. Brains were post-fixed for 2 h and cryoprotected overnight in buffer containing 20% sucrose.

Coronal sections of 10 μm thick through the hypothalamus were obtained using a cryostat (HM525 NX Epredia, Fisher Scientific), mounted on gelatine-coated slides, and processed for indirect immunohistochemistry.

### Immunohistochemistry

A sequential double-label immunohistochemical procedure was performed. Sections were first incubated overnight with a primary monoclonal antibody against neuronal nitric oxide synthase (nNOS; dilution 1:500) (Merck, Barcelona, Spain). After rinsing, sections were incubated with peroxidase-labeled goat anti-mouse F(ab’)□ fragments (1:150) (Biosys, Paris, France). Peroxidase activity was detected using 4-chloro-1-naphthol in Tris-HCl buffer (pH 7.6) with hydrogen peroxide, yielding a blue reaction product.

Subsequently, endogenous peroxidase activity was quenched using 0.3% H□O□, a standard condition that effectively suppresses residual enzymatic activity while preserving antigenicity, before incubation with the second primary antibody and visualization using a distinct chromogen. Following completion of the first immunohistochemical reaction, sections were processed for a second immunohistochemical staining using primary polyclonal antibodies against dopamine D1 receptor (D1R) (1:500) or dopamine D2 receptor (D2R) (1:500) (Abyntek, Bilbao, Spain). After rinsing, sections were incubated with peroxidase-labeled goat anti-rabbit F(ab’)□ fragments (1:150) (Biosys, Paris, France). Peroxidase activity, in this case, was detected using 3-amino-9-ethylcarbazole in acetate buffer (pH 5.0), producing a red reaction product.

### Controls

Control procedures included omission of primary antibodies and sequential processing controls to exclude cross-reactivity between detection systems. No immunoreactivity was observed in control sections.

### Quantitative analysis

Quantitative analysis was performed in thirty sections per animal along the rostrocaudal axis using a systematic sampling of the PVN. The sections were analysed and images acquired using a light microscope (DM4000B; Leica Microsystems, Wetzlar, Germany), equipped with a digital camera (LEICA DFC300 FX). nNOS-immunoreactive neurons were identified by cytoplasmic blue staining, whereas dopaminergic receptor immunoreactivity frequently appeared as a red punctate pattern associated with neuronal profiles. Only complete neuronal profiles showing clear immunoreactivity for one or both markers were included in the analysis. nNOS-ir neurons were counted in ten alternate sections using Leica Q-Win® software (Analysis Image System Leica Q-Win®, Barcelona, Spain). In the other twenty alternate sections, double labeling nNOS/D1R neurons (ten sections) and nNOS/D2R neurons (ten sections) were counted using Leica Q-Win® software. Cell counts were conducted independently by two blinded investigators.

Data are expressed as the mean number of neurons per section. The proportion of nNOS-ir neurons co-expressing D1R or D2R was calculated relative to the total nNOS-ir population.

## Results

Immunohistochemical analysis of coronal sections along the rostrocaudal axis of the PVN revealed numerous nNOS-immunoreactive (nNOS-ir) neurons distributed throughout the nucleus. These neurons showed a blue reaction product localized in the cytoplasm and proximal processes (Figure 1a).

**Figure 1.**
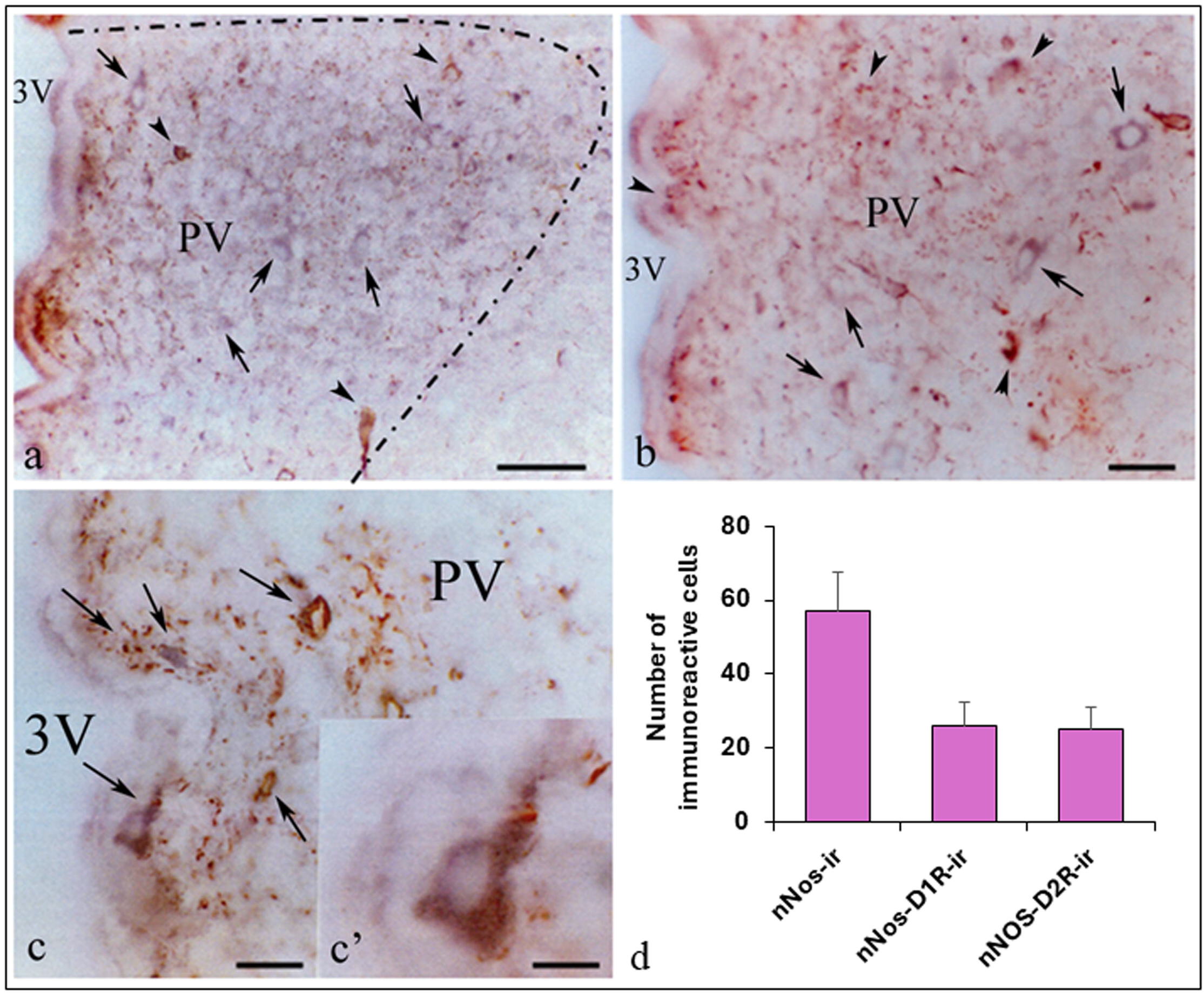
Colocalization of neuronal nitric oxide synthase (nNOS) and dopamine receptors in the paraventricular nucleus (PVN). (a) Representative coronal section showing nNOS-immunoreactive (nNOS-ir) neurons (arrows), visualized by a blue reaction product. A subset of these neurons also exhibits dopamine D2 receptor (D2R) immunoreactivity (arrowheads), detected as a red reaction product. The boundaries of the PVN are indicated by a dashed line. (b) Higher magnification of the PVN showing nNOS-ir neurons (arrows) and neurons co-expressing dopamine D1 receptor (D1R) immunoreactivity (arrowheads). (c, c′) High-magnification views illustrating nNOS-ir neurons that also display D2R immunoreactivity, indicating colocalization within the same neuronal profiles (arrows). (d) Quantitative analysis of the mean number of nNOS-ir neurons, as well as nNOS/D1R- and nNOS/D2R-immunoreactive neurons per section. Approximately 45% of nNOS-ir neurons co-express D1R or D2R. Data are expressed as mean ± SD.

Subsets of nNOS-ir neurons also exhibited immunoreactivity for dopamine D2 receptor (Figure 1a,c,c′) or dopamine D1 receptor (Figure 1b). These neurons showed a red reaction product that occupied part or almost all the cytoplasm and proximal processes. Colocalization was identified by the presence of both chromogenic reaction products within the same neuronal profiles.

Quantitative analysis yielded an average of 57.3 ± 10.4 nNOS-ir neurons per section. Of these, 26.0 ± 6.4 neurons per section co-expressed D1R immunoreactivity (≈45.4%), whereas 25.3 ± 6.0 neurons per section co-expressed D2R immunoreactivity (≈44.1%) (Figure 1d). These findings indicate that nearly half of the nitrergic neuronal population in the PVN expresses dopaminergic receptors. No significant differences were observed between the proportions of nNOS neurons co-expressing D1R and D2R.

## Discussion

The present study provides anatomical evidence for the coexistence of dopamine receptors and nNOS within neurons of the PVN and demonstrates that subsets of nNOS-ir neurons in the rat PVN also express dopamine D1 and D2 receptors. These findings support the existence of a structural substrate for a direct interaction between dopaminergic and nitrergic signaling systems within this hypothalamic nucleus, consistent with previous functional studies (Melis and Argiolas, 2011; Hull et al., 1999). Although immunofluorescence techniques are often used in colocalization studies, the sequential immunoperoxidase approach employed here offers high anatomical resolution and stable chromogenic labeling, facilitating the identification of neuronal morphology within the PVN. The use of primary antibodies raised in different species, together with sequential peroxidase inactivation, minimizes the risk of cross-reactivity and supports the reliability of the observed labeling patterns.

The punctate pattern of dopaminergic receptor immunoreactivity observed in association with nNOS-positive neurons is consistent with receptor localization at the membrane or subcellular compartments and has been described in previous studies (Missale et al., 1998).

Importantly, quantitative analysis revealed that approximately 45% of nNOS-positive neurons co-express D1R or D2R, indicating that dopaminergic modulation is restricted to a substantial but defined subpopulation of nitrergic neurons. This partial overlap suggests a selective anatomical substrate through which dopamine may regulate nitric oxide signaling within the PVN, and the similar proportions of nNOS neurons expressing D1R and D2R suggest that both receptor subtypes may contribute comparably to dopaminergic modulation within the PVN.

These observations are consistent with previous functional studies showing that dopaminergic activation in the PVN induces penile erection through nitric oxide-dependent mechanisms (Melis et al., 1996; Sanna et al., 2012; Melis et al., 2022). Moreover, pharmacological evidence indicates that modulation of dopaminergic and related signaling pathways can directly influence nitric oxide production within the PVN, affecting erectile responses (Melis et al., 2000; Melis and Argiolas, 2021). The presence of dopamine receptors in nNOS-expressing neurons supports the hypothesis that dopamine may directly modulate nitrergic neuronal activity.

The identification of both D1R- and D2R-expressing subpopulations further suggests a complex regulatory framework, as these receptor subtypes are associated with distinct intracellular signaling pathways.

Although the present study includes quantitative assessment, it remains descriptive and does not address functional activity. Therefore, further studies are required to determine the physiological relevance of these interactions. Taken together, these findings provide a structural basis for dopaminergic modulation of nitrergic signaling within the PVN and contribute to a more comprehensive understanding of the neurochemical integration underlying central mechanisms of male sexual function.

## Conclusion

The present study provides anatomical and quantitative evidence for the colocalization of dopamine D1 and D2 receptors with nNOS in neurons of the rat PVN. This organization provides a structural basis for dopaminergic modulation of nitrergic signaling and supports a framework for dopaminergic–nitrergic interaction within the PVN in the central regulation of male sexual function.

## Acknowledgements

This work was supported by the International Research Award 2000 (Abbott Laboratories Inc.).

